# Common molecular signatures between coronavirus infection and Alzheimer’s disease reveal targets for drug development

**DOI:** 10.1101/2023.06.14.544970

**Authors:** Morteza Abyadeh, Vijay K. Yadav, Alaattin Kaya

## Abstract

Cognitive decline has been reported as a common consequence of COVID-19, and studies have suggested a link between COVID-19 infection and Alzheimer’s disease (AD). However, the molecular mechanisms underlying this association remain unclear. To shed light on this link, we conducted an integrated genomic analysis using a novel Robust Rank Aggregation method to identify common transcriptional signatures of the frontal cortex, a critical area for cognitive function, between individuals with AD and COVID-19. We then performed various analyses, including the KEGG pathway, GO ontology, protein-protein interaction, hub gene, gene-miRNA, and gene-transcription factor interaction analyses to identify molecular components of biological pathways that are associated with AD in the brain also show similar changes in severe COVID-19. Our findings revealed the molecular mechanisms underpinning the association between COVID-19 infection and AD development and identified several genes, miRNAs, and TFs that may be targeted for therapeutic purposes. However, further research is needed to investigate the diagnostic and therapeutic applications of these findings.

## Introduction

The coronavirus disease 2019 (COVID-19) first appeared in early December 2019 in Wuhan, China, and in January 2020, the World Health Organization (WHO) announced a Public Health Emergency of International Concern. As of April 2023, COVID-19 had infected more than 762 million cases and took over 6.8 million lives worldwide [1]. After over three years, although extensive research has been performed to reveal mechanisms underlying COVID-19 pathogenesis, it remained poorly understood. Moreover, mounting evidence about the adverse effects of COVID-19 infection on different human organs raised concerns about the long-term sequelae of this disease and its potential role in inducing other diseases [2]. Therefore, recently, investigating the association of COVID-19 with other diseases has gained attention among researchers leading to increased reports on the role of COVID-19 in neurodegenerative diseases, diabetes, and cardiovascular diseases [2, 3]. A recent comparison of frontal cortex transcriptome data from COVID-19 patients and the elderly population showed the molecular signature of aging in the brain of COVID-19 patients [4], which suggests the potential role of COVID-19 in accelerated aging and age-related diseases. In this regard, epidemiological studies also have indicated an increased risk of developing Alzheimer’s disease (AD) following infection with COVID-19 (HR: 1.69, 95% CI: 1.53–1.72) [5]. In addition, molecular studies have shown the presence of main AD pathological hallmarks, including amyloid beta (Aβ) and phosphorylated tau protein (p-*tau)* deposition in the brain of COVID-19 infected patients [6]. AD is the most common form of dementia and a leading cause of death globally [7]. Currently, over 50 million people have dementia, and AD contributes to almost 75% of the cases, which is estimated to hit 150 million in 2050 [8]. Given the recent reports suggesting COVID-19 as the risk factor for developing AD, we should expect a higher number of AD cases than the previous estimation.

AD is a complex disease with different risk factors, typically characterized by initial memory and learning impairment followed by cognitive dysfunction, which remained incurable with only two recently approved drugs against Aβ [9, 10]. So far, a limited number of studies have investigated potential molecular mechanisms underlying the causative role of COVID-19 infection in AD development and reported possible causative mechanisms; however, molecular changes underpinning this association are still unclear [6, 11].

High-throughput technologies have proved efficient and reliable tools for comprehensively analyzing biological changes at different molecular levels [12]. Transcriptome and proteome analyses, among the most popular high-throughput approaches, significantly contributed to our understanding of disease and are now essential to biological studies. Currently, using high-throughput approaches, biological data are generated at a higher pace than interpreted. Therefore, the challenge is to extract new knowledge from existing data. A meta-analysis is a popular approach to summarize and extract the most reliable data from existing results of multiple studies, taking advantage of the increased statistical power of larger combined sample sizes [8]. Herein, in this study, we used a novel Robust Rank Aggregation (RRA) method [13, 14], which reduces errors or biases between multiple data sets to perform the meta-analysis on transcriptome datasets from AD to identify prioritized gene lists and finds commonly overlapping differentially expressed genes, (DEGs). We then compared the identified robust DEGs with transcriptome data from COVID-19 infected brains to identify common DEGs between AD and COVID-19. We further investigated protein-protein interaction and identified the hub genes within the constructed network. Subsequently, hub gene-miRNA and hub gene-transcription factor interaction network analyses have been carried out to find potential molecular targets altered commonly in both diseases.

Our analyses suggested that down-regulation of cyclic adenosine monophosphate (cAMP) signaling pathway and taurine metabolisms and up-regulation of neuroinflammatory related pathways such as Neutrophil extracellular trap (NET) formation pathway are commonly altered in AD and COVID-19 pathogenesis, and may make COVID-19 patients more susceptible to cognitive decline and AD. This information will provide a foundation for future animal and clinical studies and lead to a better understanding of the molecular mechanisms underpinning the association between COVID-19 and AD. Studies using large cohorts of COVID-19 and AD patients are needed to assess the potential therapeutic targets related to these pathways and determine the diagnostic and therapeutic potential of identified miRNA and TFs.

## Materials and Methods

### Dataset selection and processing

The raw count files of transcriptome datasets from the National Center for Biotechnology Information (NCBI) Gene Expression Omnibus (GEO) (https://www.ncbi.nlm.nih.gov/geo/ were included in our study if they met the following inclusion criteria: (1) the dataset was original; (2) reported gene expression in the same brain region of AD patients or COVID-19 infected patients; (3) both cases and controls were included. A list of differentially expressed genes (DEGs) between AD/COVID-19 cases and healthy controls were analyzed using the limma or DESeq2 R packages; Genes were considered up-regulated if fold change > 1.2 and *p*- value < 0.05 and down-regulated if fold change <0.83 and *p*-value < 0.05.

### Identfication of robuts DEGs in AD and COVID-19 datsets

Herein, we employed the Robust rank aggregation (RRA) method by using the “RobustRankAggreg” R package (Version: 1.2.1) [13], to identify robust DEGs from each dataset. To do this, the identified lists of down and up-regulated genes from each dataset were separately ranked based on their fold changes. These lists were combined into a single file, which was then subjected to the Robust RRA method. Unlike the Venn diagram analysis, which identifies shared genes, RRA identifies genes that exhibit significant fold changes across datasets, even if they are not present in all of them [13, 15]. Robust DEGs with a Bonferroni-corrected *p*-value less than 0.05 were considered statistically significant. Then, the list of AD robust DEGs was compared with the list of DEGs from COVID-19 patients using the “upset plot” to obtain common down- and up-regulated DEGs between AD and COVID-19 datasets.

### Network analysis

Protein-protein interaction (PPI) networks were analyzed using the Cytoscape-String App plugin with a confidence score > 0.05, as previously described [8]. Briefly, robust DEGs shared between AD and COVID-19 were uploaded into Cytoscape. Next, the *Homo sapiens* database in the StringDB was selected to reveal the protein interaction between differentially expressed proteins. Finally, to identify the hub genes within the protein network, CytoHubba; a plugin within Cytoscape, was utilized, and hub genes were selected based on the Maximal Clique Centrality (MCC) algorithm [16].

### Tissue-specific expression of hub genes

Genotype-Tissue Expression (GTEx) Project data, including RNA-seq data from 53 human tissue samples, was used to analyze the tissue-specific expression of the identified hub genes [17]. GTEx was accessed through the Expression Atlas database. The tissue expression levels were measured in transcripts per million (TPM), and Z-score normalization was applied to the expression levels for data visualization with a heatmap.

### Functional enrichment analysis

For functional enrichment analysis of common DEGs between AD and COVID-19 we utilized ShinyGo, a web-based tool for comprehensive gene set enrichment analysis (http://bioinformatics.sdstate.edu/go/, ShinyGO 0.77) [18]. KEGG (Kyoto Encyclopedia of Genes and Genomes) pathway, GO Biological process (BP), and GO Molecular function (MF) were then used to find the enriched terms from the submitted list of DEGs. Enriched terms with an adjusted *p*-value less than 0.05 were considered statistically significant for down and up- regulated DEGs.

### Gene–miRNA and gene-transcription factors interaction analysis

The gene–miRNA interaction analysis was carried out in the NetworkAnalyst tool [19], which uses collected data of validated miRNA-gene interaction from TarBase (which showed the complete list of miRNA for most of all DEGs) [20]. The miRNAs-DEGs network was then visualized using Cytoscape. The list of the top 5 miRNA based, on network topology measurements, including degree and betweenness centrality, were reported.

Similarly, for the gene-transcription factor (TF) interaction analysis, we employed the NetworkAnalyst tool. Official gene symbols were submitted, and related TFs were explored from ChIP-seq data, ChIP Enrichment Analysis (ChEA) (which provided the most comprehensive list of TF for all DEGs) [21]. The gene-TF interaction network was also visualized using Cytoscape.

### Literature search for the validation of the identified miRNAs and TFs via text mining

To search miRNAs and TFs from our analysis and their relation with AD or COVID-19, based on the published literature, the “batch_pubmed_dowload” function from the easyPubMed R package [22] was employed. The articles, including our identfied targets were thoroughly discussed in the discussion.

## Results

### Analysesof transcriptome data from 680 brain samples reveal data heterogeneity between AD datasets and profound transcriptomic changes in AD and COVID-19 cohorts

Three transcriptome datasets, obtained from the frontal cortex of post-mortem brains of 377 Alzheimer’s patients and 223 healthy controls (GSE118553, GSE48350, and GSE33000), and a dataset for COVID-19 (GSE188847) with 19 cases and 21 controls were included in our study (16-18, 4) (**Table 1**). DEGs in each brain region were extracted using the limma and DESeq2 R packages (*p*-value < 0.05, and log2|FC|U0.263) [23, 24]. The numbers of down- and up- regulated robust DEGs from each AD dataset were varying from 664 (down-regulated:268 and up-regulated: 396, GSE118553) to 1672 (down-regulated:1009 and up-regulated: 663, GSE48350) for the AD datasets (**Table 1**). Interestingly, analyses of the COVID-19 dataset revealed 2471 down- and 3134 up-regulated genes (**Table 1**), indicating COVID-19 infection causes profound transcriptomic changes in comparson to the AD patient cohort (**Figure 1, Supplementary file 1**). Furthermore, a comparison of DEGs between the three AD datasets showed only 1 down- and 7 up-regulated genes, shared among them (**Supplementary file 1, Supplementary figure 1**). This data indicates variations between existing AD datsets, which could be related to subtype heterogeneity across AD patient cohorts [25]. It is also possible that the difference between post-mortem sample collection times might cause variation in the transcriptome dataset, as it was shown previously [26]. Among the common genes, Rabphilin 3A (RPH3A) is a small G protein acts in the late stages of neurotransmitter exocytosis that found to be down-regulated between all three AD datasets. Down regulation of RPH3A showed to be associated with dementia severity and increased β-amyloid (Aβ) concentrations [27]. On the other side, common up-regulated genes between all three AD datasets are involved in different biological pathways mostly in immune response including, Human Leukocyte Antigen - DR Alpha (HLA-DRA), Fc Fragment of IgG Receptor IIa (FCGR2A) and Cluster of Differentiation 74 (CD74). HLA-DRA gene encodes a protein subunit of the major histocompatibility complex class II (MHC II) molecule. FCGR2A gene encodes a receptor that binds to the fragment crystallizable region (Fc region) of IgG antibodies [28]. The FCGR2A receptor is expressed on immune cells such as macrophages, neutrophils, and natural killer (NK) cells; when the receptor binds to IgG antibodies, it triggers immune effector functions such as phagocytosis and antibody-dependent cell-mediated cytotoxicity (ADCC) [29]. CD74 gene encodes a protein that plays a role in antigen processing and presentation, and it is also involved in regulating of cell proliferation and survival [30] (**Supplementary file 1**).

**Figure 1.**
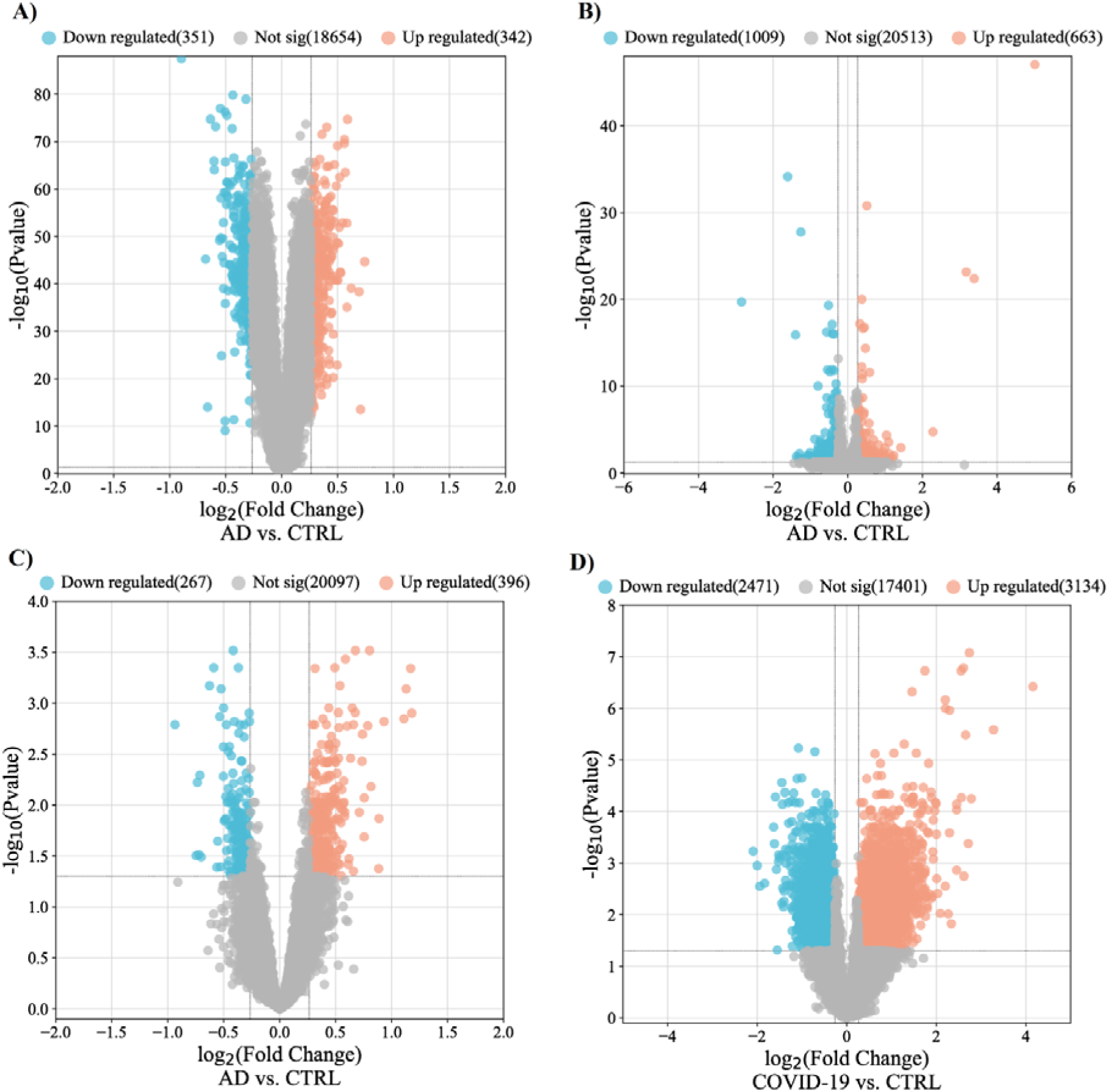
Differentially expressed genes from each datasets. Volcano plots demonstrate th thresholds for differentially expressed genes in GSE33000 (A), GSE 48350 (B), GSE118553, and GSE188847. Each data point represents a single gene. The x-axis represents the log2 fold change in expression (AD or COVID-19 vs. Control), and the −log (p-value) is plotted on the y-axis. Blue and red points indicate significantly (*p*-value < 0.05) down- and up-regulated genes with fold change lower than 0.83 and over 1.2. AD, Alzheimer’s disease; CTRL, Control; FC, Fold change.

**Table 1.** Characteristics of the selected datasets analyzed in this study.

| Datasets | Country | Number.<br>Cases/CTR<br>s | Age (yrs.)<br>AD/CTR | Post-mortem<br>interval<br>(hours)<br>AD/CTR | Disease | Number.<br>down/up-regulated<br>genes | Ref. |
| --- | --- | --- | --- | --- | --- | --- | --- |
| GSE118553 | UK | 52/27 | 82.9 ± 8.7/<br>70.6 ± 15.9 | 39.9 ± 21.3/<br>37.1 ± 20.7 | AD | 268/396 | [31] |
| GSE48350 | USA | 15/39 | 85.7 ± 6.3/<br>64.8 ± 9.5 | - | AD | 1009/663 | [32] |
| GSE33000 | USA | 310/157 | 80.6 ± 9.0/<br>63.5 ± 9.9 | 13.7 ± 7.4/<br>22.4 ± 5.8 | AD | 351/342 | [33] |
| GSE188847 | USA | 17/21 | 64.1 ± 10.5/<br>61.6 ± 14.6 | <50 | COVID-19 | 2471/3134 | [4] |

### Integrated transcriptome analysis revealed robust AD genes in COVID-19 infected brains

The RRA analysis using the integrated list of down- and up-regulated genes, including 1628 down- and 1401 up-regulated genes across AD datasets, sorted based on their fold changes yielded 44 down- and 42 up-regulated AD robust DEGs (**Figure 2 and Supplementary file 1**). The comparison between the AD robust DEGs and COVID-19 DEGs, showed 23 up and 26 down-regulated common genes (**Figure 2**). Neuronal Differentiation 6 (NEUROD6), Corticotropin Releasing Hormone (CRH), and Glutamic Acid Decarboxylase 2 (GAD2) are the top three deregulated DEGs shared between the COVID-19 dataset and robust AD genes. NEUROD6 is a basic helix-loop-helix transcription factor involved in neuronal development and differentiation, and its downregulation has been suggested as a potential biomarker for AD [34]. CHR is a gene involved in neuroendocrine responses to stress, and its level is decreased in AD patients [35]. GAD2 encodes an enzyme that catalyzes the production of gamma-aminobutyric acid from L-glutamic acid [36]. Downregulation of GAD2 and high levels of L-glutamic acid have been reported in AD patients leading to neuronal death, a phenomenon generally termed excitotoxicity [37]. While top three shared up-regulated DEGs, including Cluster of Differentiation 74 (CD74), C-X-C chemokine receptor type 4 (CXCR-4), also known as CD184, and complement component 5a receptor 1 (C5AR1) or known as CD88, are mainly involved in immune response [38]. Additionally, one DEG, Tachykinin Precursor 1 (TAC1), involved in cell-cell signaling and inflammatory response [39], was up-regulated in AD but down-regulated in COVID-19.

**Figure 2.**
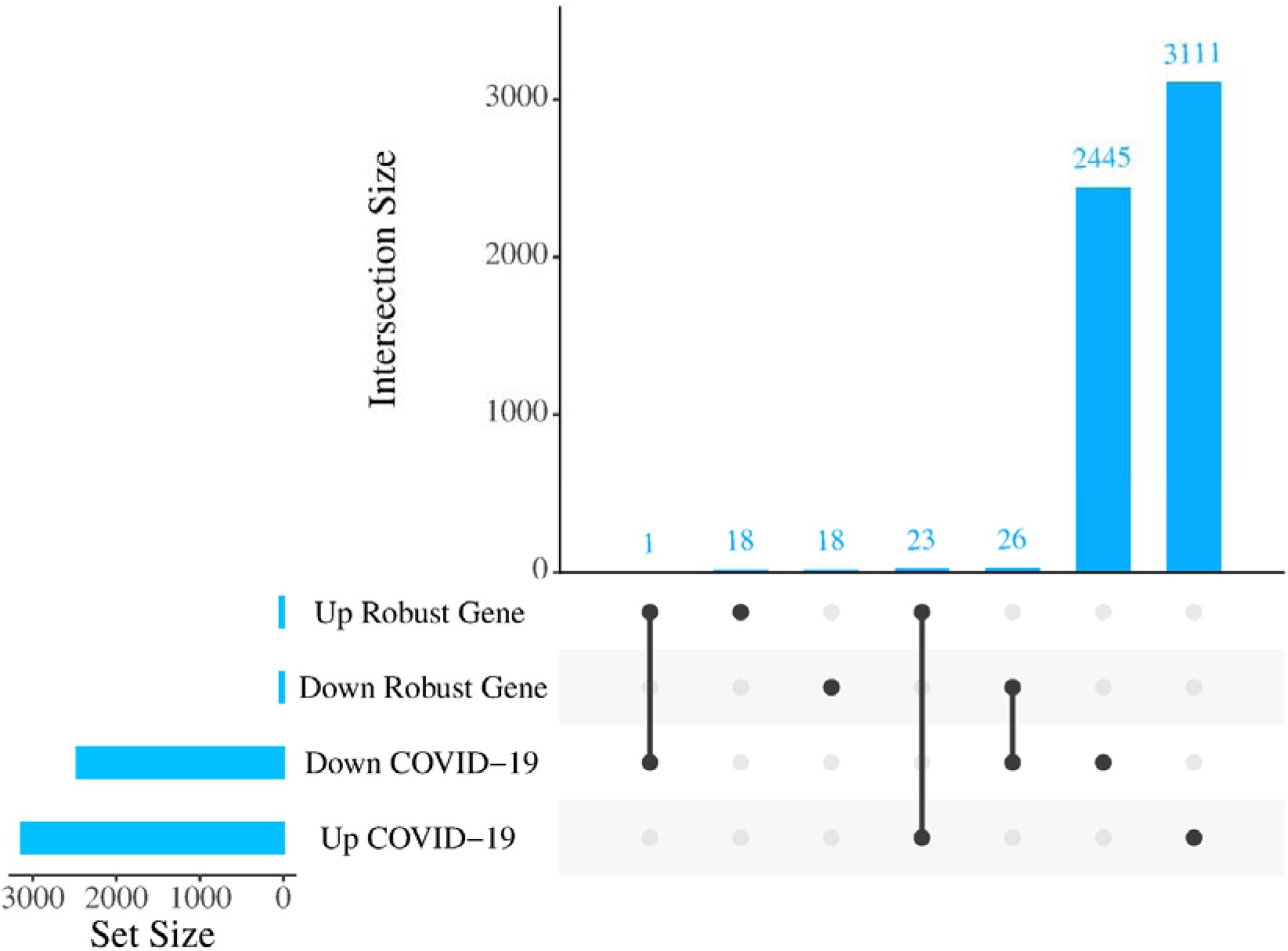
Number of commonly altered genes between AD and COVID-19. Upset plot indicating the overlap of COVID-19 DEGs with either increased or decreased expression in robust AD DEGs.

### PPI network analysis of robust DEGs identified common hub genes associated with both diseases

To understand interactions between common DEGs between AD and COVID-19 and find the hub genes within their network, we performed PPI interaction and hub gene analysis. Our analyses revealed Brain-Derived Neurotrophic Factor (BDNF), Somatostatin (SST), Glutamic Acid Decarboxylase 1/2 (GAD1/2), CRH, Vasoactive Intestinal Peptide (VIP), Glial Fibrillary Acidic Protein (GFAP), Adenylate Cyclase Activating Polypeptide 1 (ADCYAP1), CXCR-4 and NEUROD6, as the hub genes within the PPI network of common DEGs between AD and COVID-19 (**Figure 3**). In addition, our tissue-specific expression analysis further validated that these hub genes are mainly expressed in the brain, except CXCR4, a transmembrane protein of CXC chemokine receptor that showed higher expression in the blood and spleen cells (**Figure 4**, **Supplementary file 1**).

**Figure 3.**
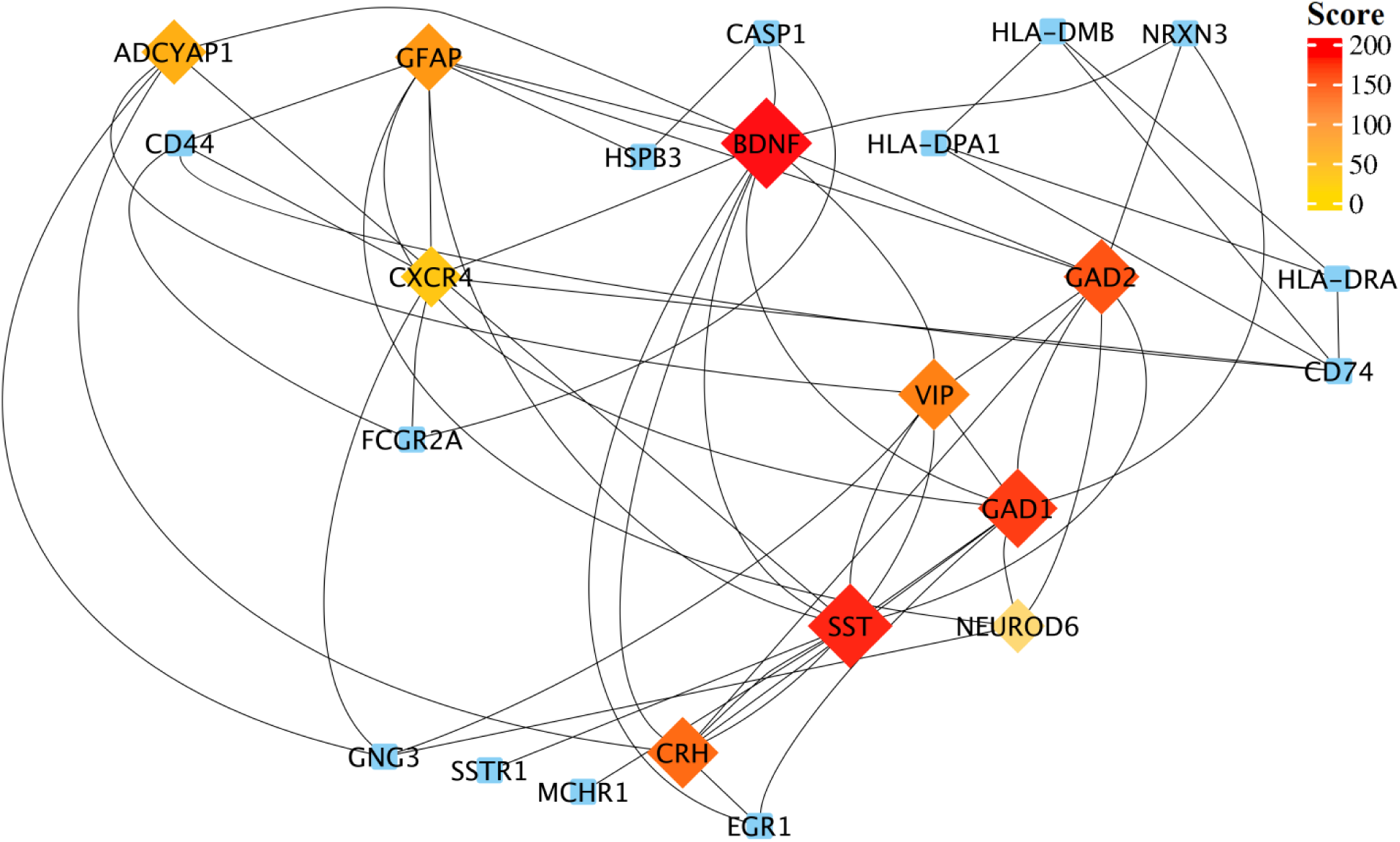
Protein-protein interaction network of those shared DEGs between robust AD and COVID-19 genes. Top-ranked hub genes are shown in rhombi shape and highlighted based on their score from the Maximal Clique Centrality (MCC) algorithm in the cytoHubba plugin of Cytoscape.

**Figure 4.**
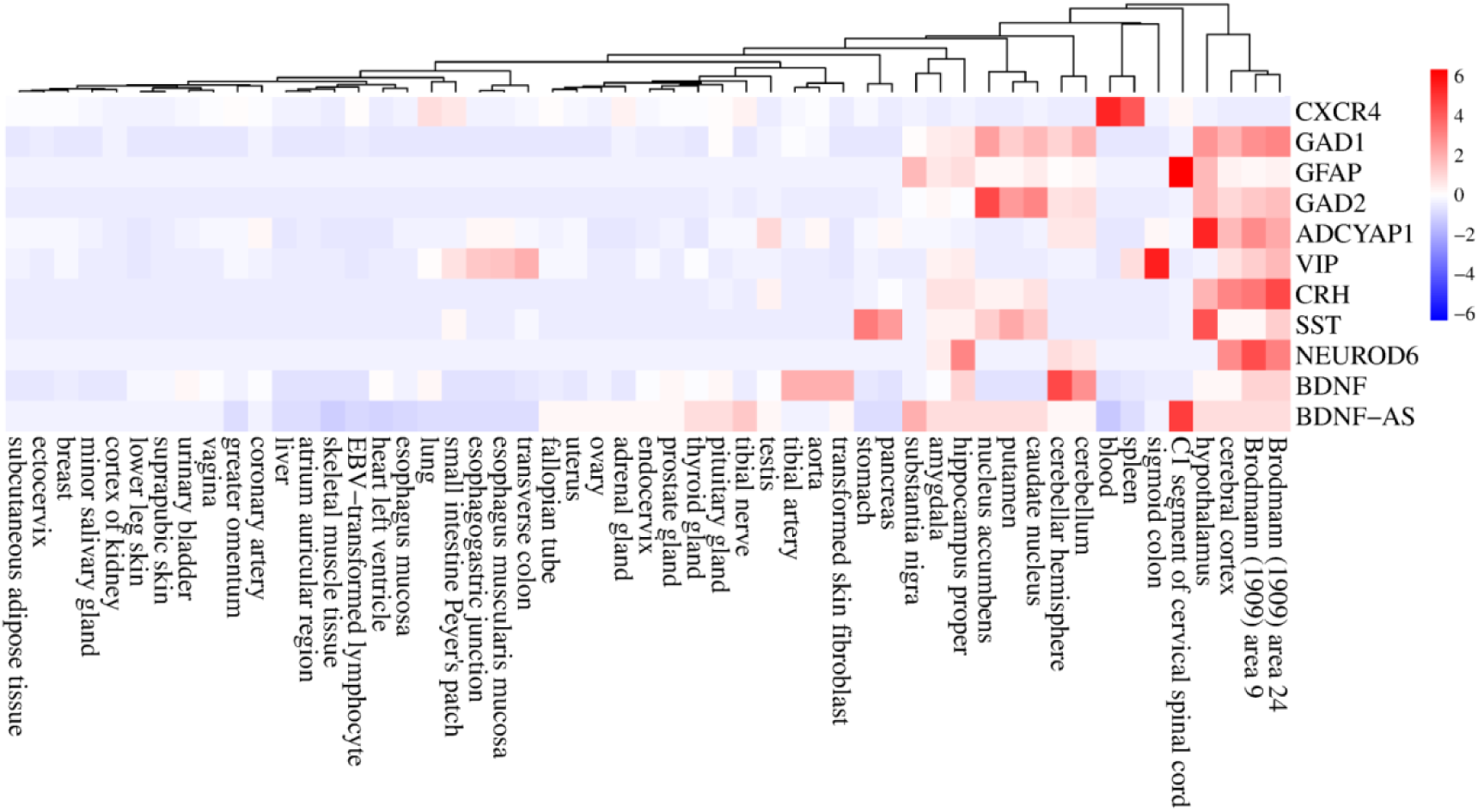
The tissue-specific expression of hub genes. The heatmap show the expression levels of the hub genes in different human tissues based on the GTEx database. Expression levels are normalized using the Z-score normalization method.

### Functional enrichment analysis revealed key pathways shared between COVID-19 and AD

We performed functional enrichment analysis to identify the common pathways altered in both AD and COVID-19. The cAMP signaling pathway was the most significant down-regulated in both diseases (Enrichment FDR= 2.24E-07) (**Figure 5**). The other significantly down-regulated enriched pathways included taurine and hypotaurine metabolism and GABAergic synapse pathway (**Figure 5**, **Supplementary file 1**). On the other hand, commonly up-regulated genes enriched in pathways involved in inflammatory responses and cell death, such as Neutrophil extracellular traps (NETs), that are structures that are formed as a defense mechanism by neutrophils to trap and neutralize invading pathogens. However, these structures can also contribute to the development of various pathophysiological conditions, including sterile inflammation and autoimmunity [40]. (**Figure 5, Supplementary file 1**). Intriguingly, pathway analysis of identified hub genes within the PPI network (**Figure 3**) also returned cAMP signaling pathway as the most commonly altered pathway (**Figure 6**), indicating a key role of this pathway in linking the COVID-19 and AD. The cAMP signaling pathway plays a role in a wide range of biological processes including metabolism, gene expression, ion transport, cell growth and differentiation, and neurotransmitter release [41]. In addition, it plays a role in the regulation of cyclic AMP response element-binding protein (CREB), a transcription factor that plays a critical role in learning and memory, as well as in neuronal development and plasticity [42].

**Figure 5.**
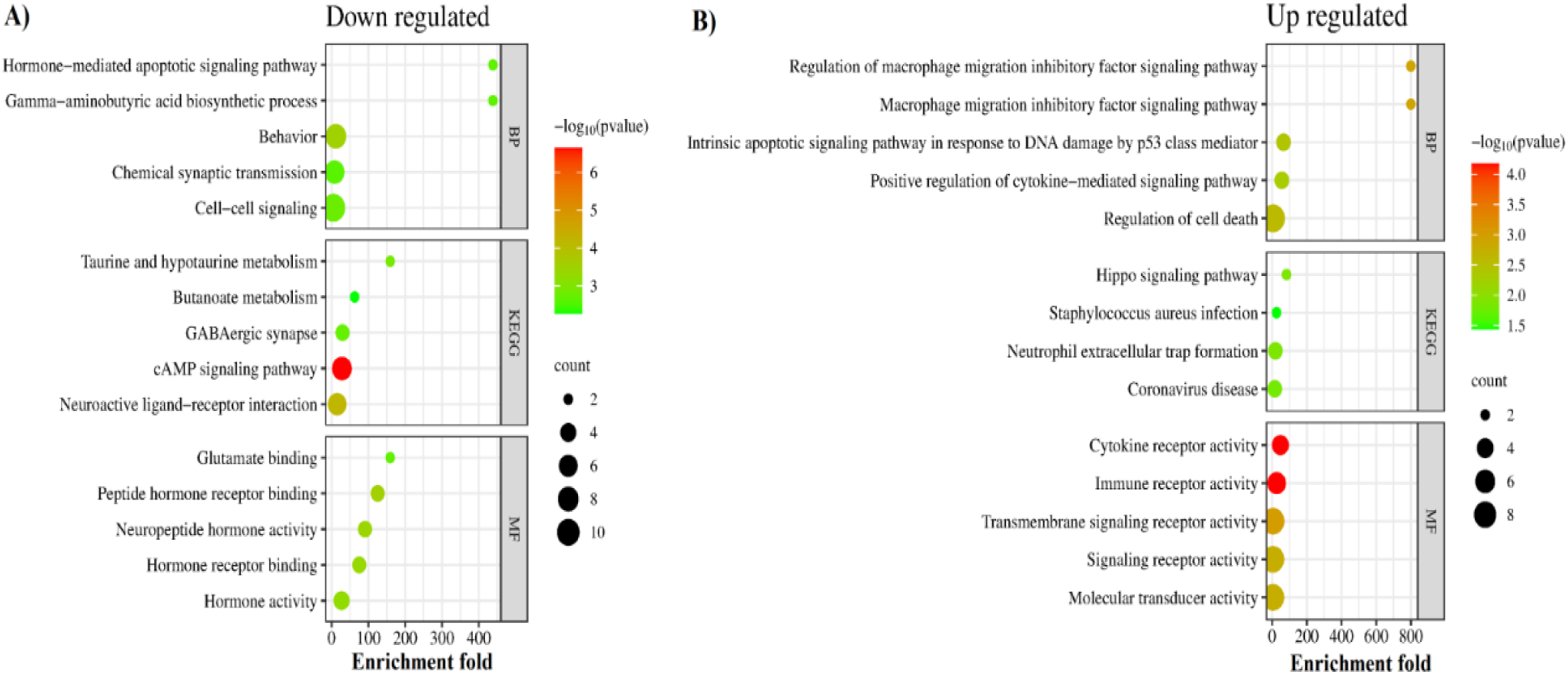
Molecular commonalities of AD and COVID-19 based on the gene expression signatures. Top 5 most significantly down- (A) and up- (B) regulated KEGG pathways, biological processes, and molecular functions enriched among the shared DEGs between robust AD and COVID-19 DEGs. The complete list of enriched terms is provided in **Supplementary file 1.**

**Figure 6.**
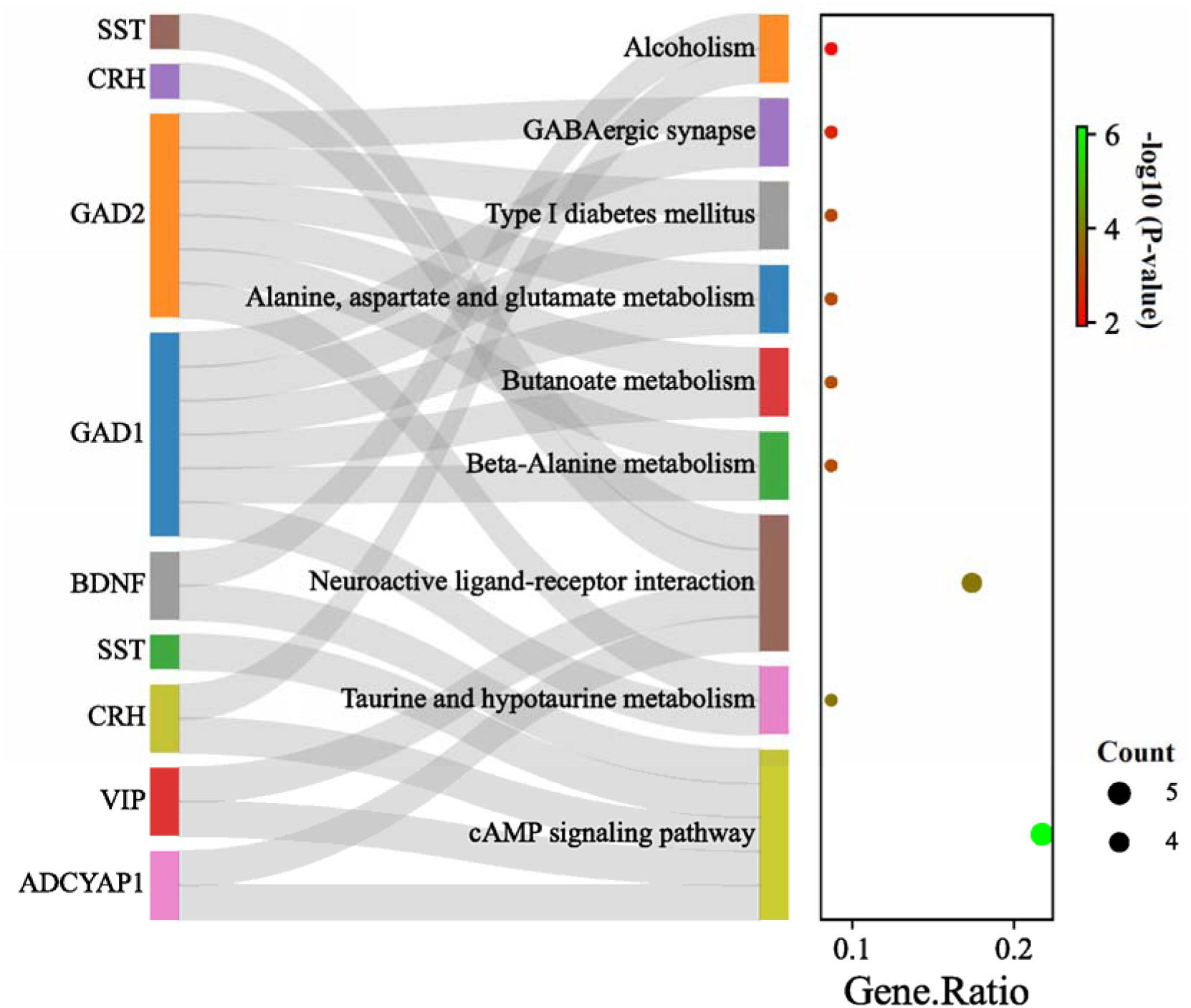
KEGG pathways enriched by identified hub genes. The left panel showing the identified hub genes, connected with their corresponding KEGG pathway. The cAMP signaling pathway is the main altered pathway including, 5 of 10 identified hub genes; however, other pathways are enriched almost by two or three genes. GAD1/GAD2 showed to be involved in most of the identified pathways. The right panel shows the gene ratio for each enriched pathway. Dot size and color indicate gene count and log10 p-values for each enriched pathway.

### Gene–miRNA and gene-TF interaction analysis revealed regulatory networks of miRNAs and TFs interacting with common hub genes

We also performed a gene-miRNA interaction analysis to identify miRNAs interacting with identified hub genes. This analysis yielded a list of 100 known and unknown miRNAs regarding their association with AD and COVID-19 (**Figure 7**). Among them, has-mir-16-5p, has-mir-27a- 3p, has-mir-130a-3p, has-mir-107, and has-mir-182-5p are identified as the top five interacting miRNAs based on their degree and betweenness within the network. Moreover, as one of the identified hub genes, BDNF was characterized by the highest number of interacting miRNAs with 35 connections. However, our analyses yielded no interacting miRNA for NEUROD6 and CRH (**Figure 7**).

**Figure 7.**
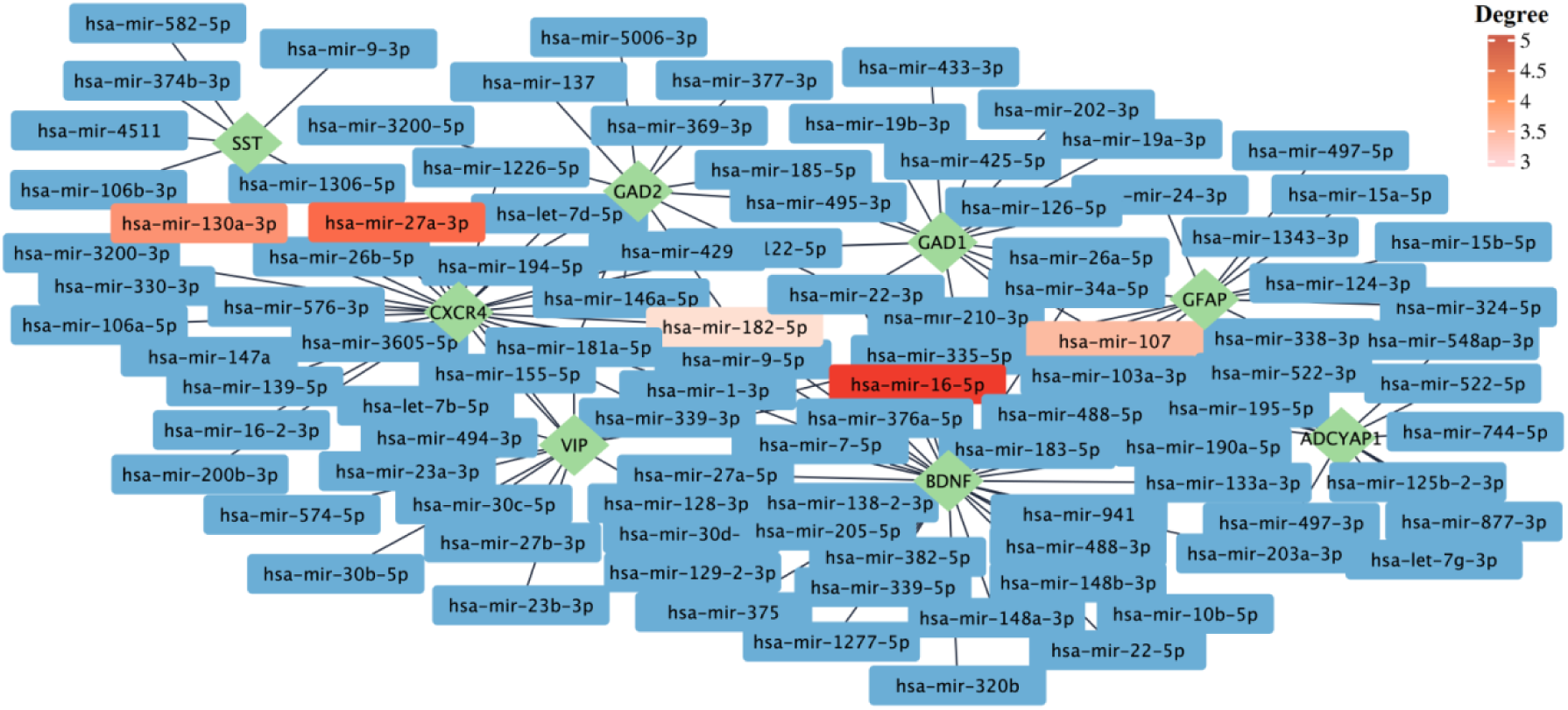
The gene–miRNA interactions network. The gene–miRNA interactions network is generated by using the TarBase database. The green rhombi show the hub genes, and the rectangles show the miRNA interacting with hub genes. The top five miRNAs based on their degrees are highlighted. The complete list of miRNAs interacting with hub genes can be found in **Supplementary file 1.**

Then, to comprehensively understand the gene regulatory network of common hub genes of COVID-19 and AD, we also performed gene-TF interaction analysis. Our analysis identified 104 known and unknown TFs related to their association with AD and COVID-19 (**Figure 8**). The top five TFs (based on network degree) include; Suppressor of Zeste 12 Protein Homolog (SUZ12), Switch-Independent 3 Family Transcription Repressor B (SIN3B), RE1-Silencing Transcription Factor (REST), Signal Transducer and Activator of Transcription 3 (STAT3) and B-Cell-Specific Moloney Murine Leukemia Virus Integration Site 1 (BMI1) (**Figure 8**).

**Figure 8.**
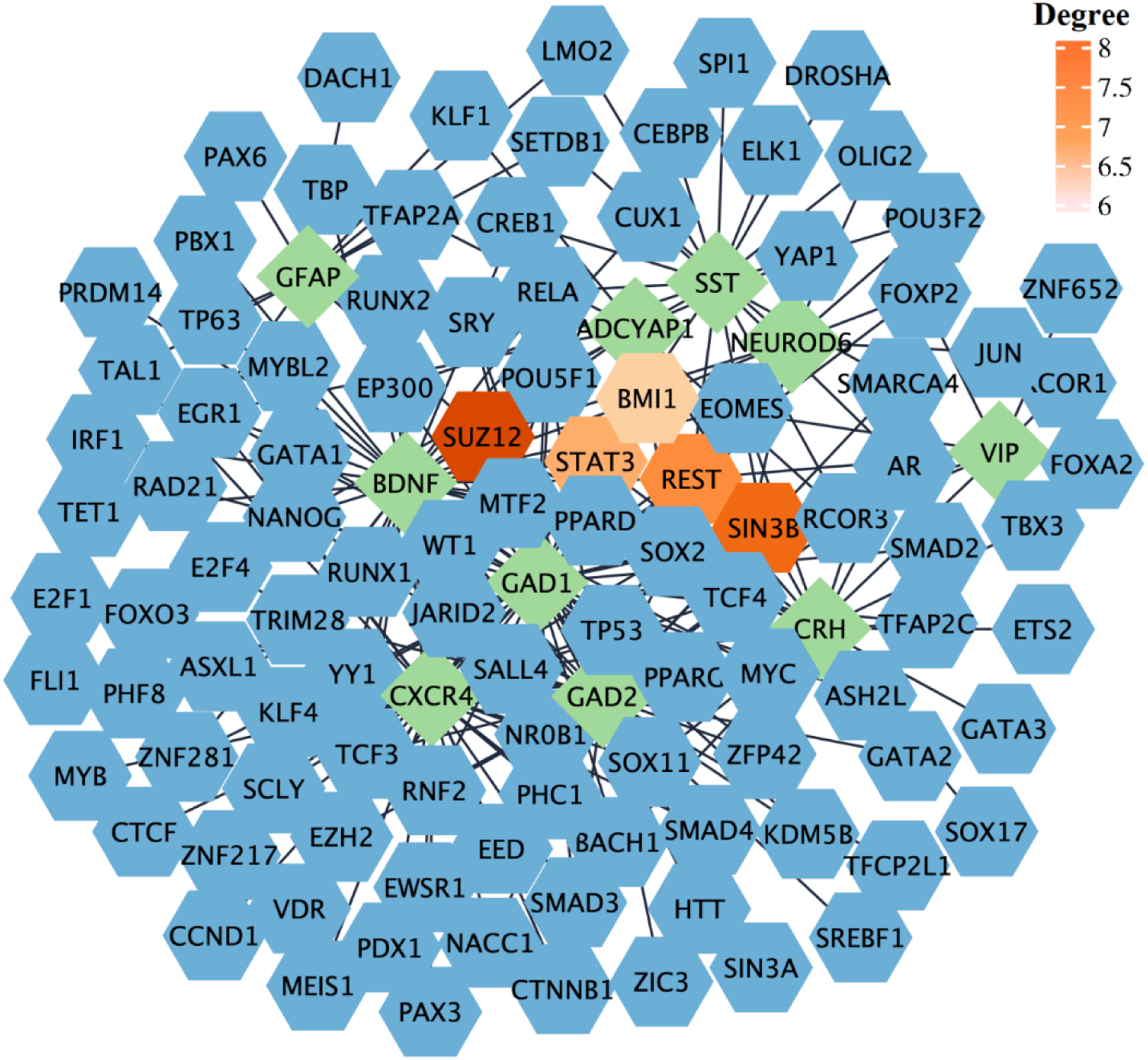
Gene-TF interaction network. The network was generated based on the ChEA database. The green rhombi show the hub genes, and the blue hexagons show the TFs interacting with hub genes, The top five identified TFs based on their degree are highlighted.

### Data mining approach for validating association of identified miRNA and TF targets in AD

Data validation using a literature mining approach was performed as a complementary analysis to our bioinformatic study to validate and interpret our results more comprehensively and accurately. Data validation was performed for the top five identified miRNAs and TFs. The results of this analysis further identified published reports on the role of some of these targets in AD and/or COVID-19 and suggested that these identified miRNAs and TFs might be utilized as pharmacological targets. In addition, our analysis also revealed no published reports for some of our identified targets. The number of published papers on the top 5 miRNAs and TFs in each disease has been shown in **Table 2** (Details are provided in **Supplementary file 1**). It is important to note that in this study, we used the names of identified miRNAs or TFs along with disease names to retrieve relevant publications on their association with COVID-19 and AD. However, it is possible that some of the identified publications may not be directly related to our study. For example, our analysis identified one related paper for has-mir-182-5p, but it focused on multiple miRNA changes in prion-infected animals and discussed the role of some of these miRNAs in AD [43]. While the involvement of has-mir-182-5p in multiple pathological conditions such as cancer, Chronic heart failure and ischemia/reperfusion (I/R) kidney injury has been widely investigated [44–46], its specific role in AD and COVID-19 remains to be investigated. However, other miRNAs provided interesting evidence in line with our results, such as mir-27a-3p. Micro-27a is located on human chromosome 19 and is processed to form miR-27a-3p [47]. miR-27a-3p is implicated in various cancer types and has recently been found to be decreased in the serum and cerebrospinal fluid (CSF) of Alzheimer’s disease (AD) patients [48]. A negative association has been observed between the level of miR-27a-3p and the degree of brain amyloid-β (Aβ) deposition, indicating its involvement in the progression of AD [48]. This miRNA has recently been found to specifically interact with one of the six regions on the viral RNA that are primarily bound by specific miRNAs, and its level has been found to be increased in hospitalized COVID-19 patients as well [49, 50]. Likewise, the top identified TFs, such as STAT3, showed the highest number of published evidence for both diseases. In contrast, we found no information regarding the role of SIN3B in either disease. STAT3 is a TF with multiple key roles in development, and its increased phosphorylation has been found in the brain of both AD animal models and patients [51]. While there is currently no evidence regarding the role of SIN3B in COVID-19 or AD, its expression has been shown to be required for cellular senescence, and its down-regulation is associated with tumor progression [52].

**Table 2.** Number of published studies on the role of identified miRNAs and TFs in AD and COVID-19

|  | Target name | Number of identified studies |  |
| --- | --- | --- | --- |
|  |  | AD | COVID-19 |
| miRNA | Has-mir-16-5p | 19 | 8 |
|  | Has-mir-27a-3p | 11 | 6 |
|  | Has-mir-103a-3p | 2 | 0 |
|  | Has-mir-107 | 49 | 1 |
|  | Has-mir-182-5p | 1 | 0 |
| TF | SUZ12 | 3 | 0 |
|  | SIN3B | 0 | 0 |
|  | REST | 38 | 1 |
|  | STAT3 | 247 | 157 |
|  | BMI1 | 10 | 0 |

## Discussion

In this study, we comprehensively analyzed transcriptome datasets of 680 brain samples, using a novel integrated genomic approach RRA method to identify robust genes altered in AD and COVID-19 infected brains. Our analysis showed that downregulation of cAMP signaling, Taurine and hypotaurine metabolism and GABAergic synapse pathways, and upregulation of inflammatory pathways such as neutrophil extracellular trap formation pathways are common signatures of both disease conditions. The cAMP signaling pathway is crucial for many biological processes. Its deregulation has been associated with aging and age-related diseases, and increases in cAMP levels have been shown to reverse some of the adverse effects of aging [53]. It has also been shown to play a key role in long-term memory formation [54]. Decreased level of cAMP has been reported in AD animal models, and pharmacological interventions that increase cAMP levels have been shown to be beneficial for neuronal protection, learning, and memory improvement in animal AD models [55, 56]. Also, the elevation of intracellular cAMP proved protective against COVID-19 Immunoglobulin G-induced procoagulant platelets and was suggested as a potential therapeutic target [57]. Intriguingly, taurine deficiency has been suggested as a critical driver of aging, and using taurine supplements increased health and life span [58]. In addition, the level of taurine has been reported to be decreased in the blood and CSF of AD patients and was associated with cognitive scores [59]. Similarly, the serum level of taurine in COVID-19-infected patients is decreased [60]. Neuroprotective effects of taurine have also been reported in animal models of AD and Parkinson’s disease (PD), reducing the Aβ aggregation and inhibiting neuroinflammation and microgliosis [59]. Intriguingly, adding taurine to drinking water significantly improved cognitive impairment and memory in mouse models of AD [61]. In contrast, there are also reports indicating increased plasma levels of taurine in both COVID-19 and AD patients [62, 63]. Increased transportation of taurine across the blood-brain barrier (BBB) has been reported during oxidative stress conditions, and NEUROD6 has been shown to be a key regulator of taurine transport through BBB [62].

On the other hand, the upregulation of inflammatory pathways, such as the neutrophil extracellular trap formation pathway, has been reported to be involved in AD [64]. The role of microglia, as the brain’s resident population of immune cells in neuroinflammation and AD has been widely investigated. However, it has been recently suggested that infiltrating blood-derived neutrophils into the central nervous system (CNS) can also contribute to AD pathogenesis and cognitive impairment [65]. In addition, neutrophils enter the CNS before the onset of cognitive impairment and are found to be highly abundant when memory loss is first observed. Blocking this process might have therapeutic potential to restore cognitive function [65]. Neutrophils are potent sources of reactive oxygen species (ROS), and their activation and associated oxidative stress have been shown to be associated with AD pathology in humans, and neutrophil-related inflammatory factors have been suggested as the potential biomarkers to predict memory and executive function decline in patients with mild AD [66]. In both human and animal models of AD, neutrophils are found to be co-localized with senile plaques and stained for NET markers (27). In addition, neutrophil adhesion in brain capillaries decreased cortical blood flow, leading to memory impairment in the mouse AD model [67]. Furthermore, neutrophils play a key role in pathogen clearance through phagocytosis, NETs, and generating ROS [68]. SARS-CoV-2 infected human and animal models showed an increased number of neutrophils and proteins associated with neutrophil degranulation [69, 70], and neutrophil degranulation has been identified as one of the main enriched pathways in proteomics analysis of COVID-19 patients [71]. Altogether, the functional enrichment analysis results showed that the downregulation of biological functions related to anti-inflammatory responses and, accordingly, upregulation of inflammatory responses may, in part, contribute to post-COVID-19 cognitive impairment and possibly AD development.

The result of hub gene analysis also showed some of the well-known genes involved in signal transduction, memory, and cognition, such as BDNF, a key neuroplasticity regulator. Its downregulation has been reported as one of the primary mediators of AD, multiple sclerosis (MS), and PD pathogenesis [72]. Interestingly BDNF also protects neurons against hypoxia and inflammation-induced pathogenesis, the key pathological events in COVID-19 [73, 74]. Recent studies reported lower serum levels of BDNF in COVID-19 infected patients and indicated a direct association between BDNF and cognitive decline in COVID-19 patients [75]. SST, another identified hub gene, is a multi-functional neuropeptide in a subpopulation of GABAergic interneurons [76]. SST’s expression level is shown to decrease with age and contributes to the formation of Aβ plaque deposition [76, 77]. While no data is available about Somatostatin in COVID-19 patients, its analogs are suggested as potential drugs for treating respiratory failure in diseases like COVID-19 [78]. Most of the identified hub genes are retrieved from down- regulated genes; however, there are two hub genes from up-regulated genes, including GFAP and CXCR4. GFAP, an astrocytic cytoskeletal protein, was up-regulated in AD patients and cognitively normal older adults at risk of AD and correlated with amyloid-PET positivity and worse outcomes in global cognition [79, 80]. Increased plasma level of GFAP is also suggested as a potential prognostic marker in COVID-19, associated with mortality risk [81]. CXCR4 is another up-regulated hub gene in AD and COVID-19 patients [82, 83]. CXCR4 is a G proteinUcoupled receptor that binds to CXCL12 and triggers downstream signaling pathways associated with inflammatory pathways [82]. CXCR4 is involved in neuronal guidance and apoptosis via astroglial signaling and microglial activation [84]. Aβ plaques have an attraction effect on microglia, leading to the activation of an inflammatory cascade. This cascade involves the stimulation of CXCR4-dependent signaling by CXCL12 in both microglia and astrocytes, resulting in the release of pro-inflammatory cytokines, including tumor necrosis factor α (TNF-) [85].

Next, we analyzed the miRNA-gene interaction network to identify miRNAs interacting with hub genes. During the last decade, exploring the involvement of miRNAs in various human diseases has gained much attention, mainly due to their crucial function in gene regulation, through mediating mRNA degradation and regulating transcription and translation via both canonical and canonical and non-canonical mechanisms [86]. Our analysis returned 100 miRNAs; among them, has-mir-16-5p was the most significant (Degree: 5; Betweenness: 1132.11) node, interacting with hub genes. Others include; miRNA has-mir-27a-3p, has-mir- 130a-3p, has-mir-107, and has-mir-182-5p. Hitherto, results about the role of mir-16-5p in AD are inconsistent. While some indicated that upregulation of this mir-16-5p by Aβ deposition can lead to neuronal cell apoptosis through targeting BCL-2 [87], others suggested a protective role of this miRNA against Aβ-induced injury by targeting BACE1 [88]. The expression level of hsa- miR-16-5p was shown to be lower in COVID-19 patients than in healthy controls [50]. This miRNA can target the many identified DEGs in macrophages (n = 15) and T cells (n = 10) in COVID-19 infected patients [89]. hsa-mir-16-5p was found to affect T cells’ cell cycle, survival, and differentiation and modulate inflammatory signaling and cytokines, including IL-1β, IL-6 and TNF-α, and NF-κB mTOR-related pathways and genes [90]. In addition, a recent study of the *in silico* analysis suggested a regulatory role of has-mir-16-5p and has-mir-27a-3p in the Angiotensin-Converting Enzyme 2 (ACE2) network [90]. The ACE2 receptor, found in several human organs, is the entry point for SARS-CoV-2 and SARS-CoV into host cells [91]. A lower level of mir-27a-3p has been reported in AD patients’ cerebrospinal fluid CSF, accompanied by high tau levels and low levels of β-amyloid [92]. It has been reported that miR-27a-3p can down-regulate Glycogen Synthase Kinase 3 Beta (GSK3β) and activate the Wnt/β-catenin signaling pathway, which ultimately helps in maintaining the integrity of the blood-brain barrier [93].

Although, in our analysis, we have only performed data mining for the top five identified miRNAs, there is also evidence of the association of several other enriched miRNAs with AD and COVID-19. For example, down-regulation of mir-124-3p, one of the mir-targeting identified hub genes, has been reported in AD patients [94]. Interestingly, mir-124-3p showed to decrease abnormal hyperphosphorylation of tau protein and subsequent cell apoptosis through regulating Caveolin-1, phosphoinositide 3-kinase (PI3K), phospho-Akt (Akt-Ser473)/Akt, phospho-GSK- 3β (GSK-3β-Ser9)/GSK-3β pathway [95]. Conversely, upregulation of mir-7-5p has been shown in AD patients, contributing to AKT and GSK3β dephosphorylation and insulin resistance in neuronal cells, accelerating the progression of Aβ plaque and neurofibrillary tangles (NFTs) formation via multiple mechanisms [96, 97]. Expression levels of both mir-124-3p and mir-7-5p showed to decrease in COVID-19 patients compared to healthy controls [98]. Intriguingly, mir- 7-5p and mir-24-3p were found to directly inhibit S protein expression and SARS-CoV-2 replication [99].

In addition, there are evidences on some of the identified TFs such as REST, and SUZ12. Consistent with the literature, our analyses revealed that while the expression of REST decreased, SUZ12 expression is increased in AD brains in comparison to the control [100, 101]. REST has been shown to protect against AD via downregulating genes that promote cell death and AD pathology and trigger stress response gene expression. In addition, REST also protects neurons against amyloid-β induced toxicity and oxidative stress, and its deficiency leads to age-related neurodegeneration [100]. The protective role of glial REST against neurodegenerative diseases also has been linked to inhibitory effects on innate immunity and inflammation [100].

STAT3 is another top-identified TF involved in various physiological processes, including immune reactions [102]. Increased phosphorylation of STAT3 and its abnormal activation has been reported in both the hippocampus of AD mouse models and post-mortem AD brain, which is critical for the secretion of cytokines involved in neuroinflammation, and correlated with the presence of reactive astrocytes in animal models of AD [103]. Additionally, STAT3 could serve as a transcriptional regulator for BACE1, the principal enzyme involved in the production of amyloid β (Aβ), and STAT3 inhibition was shown to reduce the level of BACE1 and neuroinflammation [51]. The STAT-3 inhibition as a downstream element in the IL-6/JAK/STAT-3 axis is also suggested as a therapeutic strategy to mitigate COVID-19 severity [104]. STAT-3 may play multiple roles during COVID-19 infection, such as instigating pro-inflammatory reactions, initiating the cytokine storm, disrupting the immune response balance, impairing anti-viral immune responses, and intensifying lymphopenia [104, 105].

Overall, our data conclude that our gene regulatory network analysis identified known and unknown genetic components of biological pathways associated with COVID-19 infection and AD development and might be targeted for therapeutic purposes to reduce the risk or delay the development of COVID-19-related neurological pathologies.

In conclusion, our study has unveiled significant insights into the molecular mechanisms linking COVID-19 infection and AD development. Through comprehensive transcriptomic analysis, we have identified shared transcriptional signatures in the frontal cortex of individuals affected by AD and COVID-19, thereby illuminating the common pathways and biological processes involved. Notably, we have observed downregulation of the cAMP signaling pathway and perturbations in taurine metabolisms, alongside upregulation of neuroinflammatory pathways such as NET formation. These findings suggest that the convergence of these molecular components may render COVID-19 patients more vulnerable to cognitive decline and AD. Moreover, our study has pinpointed several promising therapeutic targets for COVID-19 and AD, encompassing genes, miRNAs, and TFs. Restoration of cAMP levels and supplementation of taurine hold potential as neuroprotective interventions.

Modulating inflammatory responses and targeting specific hub genes, miRNAs, and TFs also present prospects for mitigating cognitive impairment in individuals experiencing post-COVID-19 complications and AD progression. Nevertheless, it is crucial to note that the therapeutic targets identified in our study necessitate rigorous experimental validation to ascertain their clinical efficacy and safety. Therefore, further investigation employing animal models and large-scale clinical cohorts is imperative to validate and expand our findings.

## Acknowledgment

Not applicable.

## Data Availability

The data supporting the findings of this study are available within the article and/or its supplementary material.

## Abbreviations

COVID-19: Coronavirus disease 2019
AD: Alzheimer’s disease
WHO: World Health Organization
P-tau: Phosphorylated tau protein (p-tau)
RRA: Robust Rank Aggregation
MF: Molecular function
BP: Biological process
DEG: Differentially expressed gene
cAMP: Cyclic adenosine monophosphate
MCC: Maximal Clique Centrality
GTEx: Genotype-Tissue Expression
TPM: Transcripts per million
CSF: Cerebrospinal fluid
CNS: Central nervous system
ROS: Reactive oxygen species
PPI: Protein-protein interaction
TF: Transcription factor
BDNF: Brain-derived neurotrophic factor
SST: Somatostatin
GAD1/2: Glutamic acid decarboxylase ½
CRH: Corticotropin Releasing Hormone
VIP: Vasoactive Intestinal Peptide
GFAP: Glial fibrillary acidic protein
ADCYAP1: Adenylate Cyclase Activating Polypeptide 1
CXCR-4: C-X-C chemokine receptor type 4
NEUROD6: Neuronal Differentiation 6
NFT: Neurofibrillary tangles
NET: Neutrophil extracellular trap

---

**Supplementary Figure 1:**
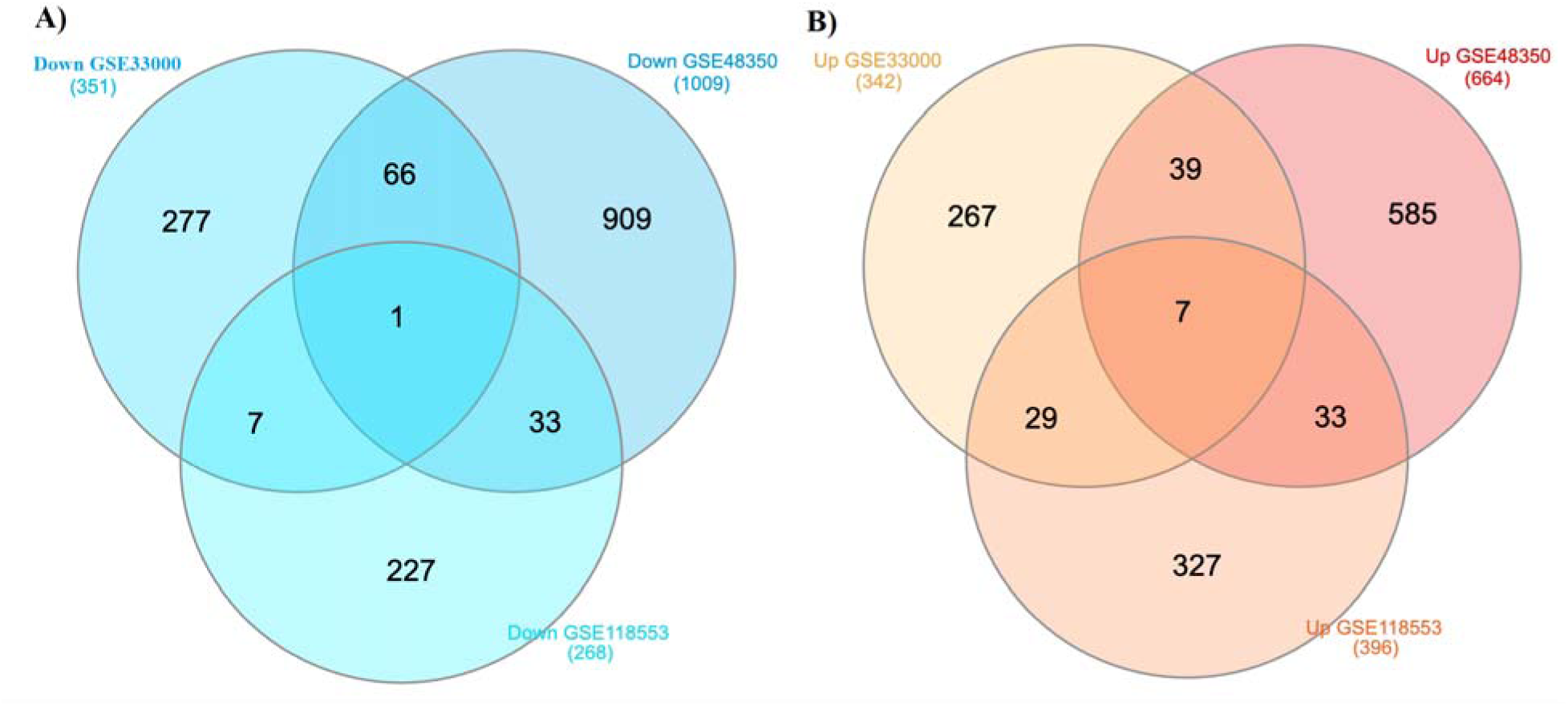
Comparison of the three transcriptome datasets of AD. (A) Commonly down- and (B) up-regulated genes are shown by the Venn diagram.

## References

[1] Organization WH (2021).

[2] Davis HE, McCorkell L, Vogel JM, Topol EJ (2023) Long COVID: major findings, mechanisms and recommendations. Nature Reviews Microbiology, 1-14.

[3] Arbov E, Tayara A, Wu S, Rich TC, Wagener BM (2022) COVID-19 and Long-Term Outcomes: Lessons from Other Critical Care Illnesses and Potential Mechanisms. American Journal of Respiratory Cell and Molecular Biology 67, 275–283.

[4] Mavrikaki M, Lee JD, Solomon IH, Slack FJ (2022) Severe COVID-19 is associated with molecular signatures of aging in the human brain. Nature Aging, 1-8.

[5] Wang L, Davis PB, Volkow ND, Berger NA, Kaelber DC, Xu R (2022) Association of COVID- 19 with new-onset Alzheimer’s disease. Journal of Alzheimer’s Disease, 1-4.

[6] Reiken S, Sittenfeld L, Dridi H, Liu Y, Liu X, Marks AR (2022) Alzheimer’s-like signaling in brains of COVID-19 patients. Alzheimer’s & Dementia 18, 955–965.

[7] Chitranshi N, Kumar A, Sheriff S, Gupta V, Godinez A, Saks D, Sarkar S, Shen T, Mirzaei M, Basavarajappa D (2021) Identification of novel cathepsin B inhibitors with implications in Alzheimer’s disease: Computational refining and biochemical evaluation. Cells 10, 1946.

[8] Abyadeh M, Tofigh N, Hosseinian S, Hasan M, Amirkhani A, Fitzhenry MJ, Gupta V, Chitranshi N, Salekdeh GH, Haynes PA (2022) Key genes and biochemical networks in various brain regions affected in Alzheimer’s disease. Cells 11, 987.

[9] Cummings J (2023) Anti-Amyloid Monoclonal Antibodies are Transformative Treatments that Redefine Alzheimer’s Disease Therapeutics. Drugs, 1-8.

[10] Abyadeh M, Gupta V, Gupta V, Chitranshi N, Wu Y, Amirkhani A, Meyfour A, Sheriff S, Shen T, Dhiman K (2021) Comparative analysis of aducanumab, zagotenemab and pioglitazone as targeted treatment strategies for Alzheimer’s disease. Aging and disease 12, 1964.

[11] Green R, Mayilsamy K, McGill AR, Martinez TE, Chandran B, Blair LJ, Bickford PC, Mohapatra SS, Mohapatra S (2022) SARS-CoV-2 infection increases the gene expression profile for Alzheimer’s disease risk. Molecular Therapy-Methods & Clinical Development 27, 217–229.

[12] Abyadeh M, Gupta V, Chitranshi N, Gupta V, Wu Y, Saks D, Wander Wall R, Fitzhenry MJ, Basavarajappa D, You Y (2021) Mitochondrial dysfunction in Alzheimer’s disease-a proteomics perspective. Expert review of Proteomics 18, 295–304.

[13] Kolde R, Laur S, Adler P, Vilo J (2012) Robust rank aggregation for gene list integration and meta-analysis. Bioinformatics 28, 573–580.

[14] Ma X, Mo C, Huang L, Cao P, Shen L, Gui C (2021) Robust Rank Aggregation and Least Absolute Shrinkage and Selection Operator Analysis of Novel Gene Signatures in Dilated Cardiomyopathy. Frontiers in Cardiovascular Medicine, 1854.

[15] Hosseinian S, Arefian E, Rakhsh-Khorshid H, Eivani M, Rezayof A, Pezeshk H, Marashi S-A (2020) A meta-analysis of gene expression data highlights synaptic dysfunction in the hippocampus of brains with Alzheimer’s disease. Scientific reports 10, 1–9.

[16] Chin C-H, Chen S-H, Wu H-H, Ho C-W, Ko M-T, Lin C-Y (2014) cytoHubba: identifying hub objects and sub-networks from complex interactome. BMC systems biology 8, 1–7.

[17] Consortium G, Ardlie KG, Deluca DS, Segrè AV, Sullivan TJ, Young TR, Gelfand ET, Trowbridge CA, Maller JB, Tukiainen T (2015) The Genotype-Tissue Expression (GTEx) pilot analysis: multitissue gene regulation in humans. Science 348, 648–660.

[18] Ge SX, Jung D, Yao R (2020) ShinyGO: a graphical gene-set enrichment tool for animals and plants. Bioinformatics 36, 2628–2629.

[19] Zhou G, Soufan O, Ewald J, Hancock RE, Basu N, Xia J (2019) NetworkAnalyst 3.0: a visual analytics platform for comprehensive gene expression profiling and meta-analysis. Nucleic acids research 47, W234–W241.

[20] Karagkouni D, Paraskevopoulou MD, Chatzopoulos S, Vlachos IS, Tastsoglou S, Kanellos I, Papadimitriou D, Kavakiotis I, Maniou S, Skoufos G (2018) DIANA-TarBase v8: a decade-long collection of experimentally supported miRNA–gene interactions. Nucleic acids research 46, D239–D245.

[21] Lachmann A, Xu H, Krishnan J, Berger SI, Mazloom AR, Ma’ayan A (2010) ChEA: transcription factor regulation inferred from integrating genome-wide ChIP-X experiments. Bioinformatics 26, 2438–2444.

[22] Fantini D (2019) easyPubMed: Search and retrieve scientific publication records from PubMed. R package version 2.

[23] Ritchie ME, Phipson B, Wu D, Hu Y, Law CW, Shi W, Smyth GK (2015) limma powers differential expression analyses for RNA-sequencing and microarray studies. Nucleic acids research 43, e47–e47.

[24] Love M, Anders S, Huber W (2014) Differential analysis of count data–the DESeq2 package. Genome Biol 15, 10–1186.

[25] Neff RA, Wang M, Vatansever S, Guo L, Ming C, Wang Q, Wang E, Horgusluoglu-Moloch E, Song W-m, Li A (2021) Molecular subtyping of Alzheimer’s disease using RNA sequencing data reveals novel mechanisms and targets. Science advances 7, eabb5398.

[26] Liharska LE, Park YJ, Ziafat K, Wilkins L, Silk H, Linares LM, Vornholt E, Sullivan B, Cohen V, Kota P (2023) A study of gene expression in the living human brain. medRxiv, 2023.2004. 2021.23288916.

[27] Tan MG, Lee C, Lee JH, Francis PT, Williams RJ, Ramírez MJ, Chen CP, Wong PT-H, Lai MK (2014) Decreased rabphilin 3A immunoreactivity in Alzheimer’s disease is associated with Aβ burden. Neurochemistry international 64, 29–36.

[28] Li M, Jin X, Li H, Wu G, Wang S, Yang C, Deng S (2020) Key genes with prognostic values in suppression of osteosarcoma metastasis using comprehensive analysis. BMC cancer 20, 1–13.

[29] Li X, Ptacek TS, Brown EE, Edberg JC (2009) Fcγ receptors: structure, function and role as genetic risk factors in SLE. Genes & Immunity 10, 380–389.

[30] Gold DV, Stein R, Burton J, Goldenberg DM (2011) Enhanced expression of CD74 in gastrointestinal cancers and benign tissues. International journal of clinical and experimental pathology 4, 1.

[31] Patel H, Hodges AK, Curtis C, Lee SH, Troakes C, Dobson RJ, Newhouse SJ (2019) Transcriptomic analysis of probable asymptomatic and symptomatic alzheimer brains. Brain, behavior, and immunity 80, 644–656.

[32] Berchtold NC, Coleman PD, Cribbs DH, Rogers J, Gillen DL, Cotman CW (2013) Synaptic genes are extensively downregulated across multiple brain regions in normal human aging and Alzheimer’s disease. Neurobiology of aging 34, 1653–1661.

[33] Narayanan M, Huynh JL, Wang K, Yang X, Yoo S, McElwee J, Zhang B, Zhang C, Lamb JR, Xie T (2014) Common dysregulation network in the human prefrontal cortex underlies two neurodegenerative diseases. Molecular systems biology 10, 743.

[34] Satoh J-i, Yamamoto Y, Asahina N, Kitano S, Kino Y (2014) RNA-Seq data mining: downregulation of NeuroD6 serves as a possible biomarker for alzheimer’s disease brains. Disease markers 2014.

[35] May C, Rapoport S, Tomai T, Chrousos G, Gold P (1987) Cerebrospinal fluid concentrations of corticotropin-releasing hormone (CRH) and corticotropin (ACTH) are reduced in patients with Alzheimer’s disease. Neurology 37, 535–535.

[36] Gleichmann M, Zhang Y, Wood III WH, Becker KG, Mughal MR, Pazin MJ, van Praag H, Kobilo T, Zonderman AB, Troncoso JC (2012) Molecular changes in brain aging and Alzheimer’s disease are mirrored in experimentally silenced cortical neuron networks. Neurobiology of aging 33, 205. e201–205. e218.

[37] Madeira C, Vargas-Lopes C, Brandão CO, Reis T, Laks J, Panizzutti R, Ferreira ST (2018) Elevated glutamate and glutamine levels in the cerebrospinal fluid of patients with probable Alzheimer’s disease and depression. Frontiers in psychiatry 9, 561.

[38] Engel P, Boumsell L, Balderas R, Bensussan A, Gattei V, Horejsi V, Jin B-Q, Malavasi F, Mortari F, Schwartz-Albiez R (2015) CD nomenclature 2015: human leukocyte differentiation antigen workshops as a driving force in immunology. The Journal of Immunology 195, 4555–4563.

[39] Borbely E, Hajna Z, Sandor K, Kereskai L, Toth I, Pinter E, Nagy P, Szolcsanyi J, Quinn J, Zimmer A (2013) Role of tachykinin 1 and 4 gene-derived neuropeptides and the neurokinin 1 receptor in adjuvant-induced chronic arthritis of the mouse. PloS one 8, e61684.

[40] Marković DC, Maslovarić IS, Kovačić M, Vignjević Petrinović S, Ilić VL (2023) Putative Role of Neutrophil Extracellular Trap Formation in Chronic Myeloproliferative Neoplasms. International Journal of Molecular Sciences 24, 4497.

[41] Yan K, Gao LN, Cui YL, Zhang Y, Zhou X (2016) The cyclic AMP signaling pathway: Exploring targets for successful drug discovery. Molecular medicine reports 13, 3715–3723.

[42] Wang H, Xu J, Lazarovici P, Quirion R, Zheng W (2018) cAMP response element-binding protein (CREB): a possible signaling molecule link in the pathophysiology of schizophrenia. Frontiers in molecular neuroscience 11, 255.

[43] Boese AS, Saba R, Campbell K, Majer A, Medina S, Burton L, Booth TF, Chong P, Westmacott G, Dutta SM (2016) MicroRNA abundance is altered in synaptoneurosomes during prion disease. Molecular and Cellular Neuroscience 71, 13–24.

[44] Cao M-Q, You A-B, Zhu X-D, Zhang W, Zhang Y-Y, Zhang S-Z, Zhang K-w, Cai H, Shi W-K, Li X-L (2018) miR-182-5p promotes hepatocellular carcinoma progression by repressing FOXO3a. Journal of hematology & oncology 11, 1–12.

[45] Ding C, Ding X, Zheng J, Wang B, Li Y, Xiang H, Dou M, Qiao Y, Tian P, Xue W (2020) miR- 182-5p and miR-378a-3p regulate ferroptosis in I/R-induced renal injury. Cell death & disease 11, 929.

[46] Fang F, Zhang X, Li B, Gan S (2022) miR-182-5p combined with brain-derived neurotrophic factor assists the diagnosis of chronic heart failure and predicts a poor prognosis. Journal of Cardiothoracic Surgery 17, 88.

[47] Li X, Xu M, Ding L, Tang J (2019) MiR-27a: a novel biomarker and potential therapeutic target in tumors. Journal of Cancer 10, 2836.

[48] He L, Chen Z, Wang J, Feng H (2022) Expression relationship and significance of NEAT1 and miR-27a-3p in serum and cerebrospinal fluid of patients with Alzheimer’s disease. BMC neurology 22, 1–8.

[49] Fossat N, Lundsgaard EA, Costa R, Rivera-Rangel LR, Nielsen L, Mikkelsen LS, Ramirez S, Bukh J, Scheel TK (2023) Identification of the viral and cellular microRNA interactomes during SARS-CoV-2 infection. Cell reports 42.

[50] Eyileten C, Wicik Z, Simões SN, Martins-Jr DC, Klos K, Wlodarczyk W, Assinger A, Soldacki D, Chcialowski A, Siller-Matula JM (2022) Thrombosis-related circulating miR-16-5p is associated with disease severity in patients hospitalised for COVID-19. RNA biology 19, 963–979.

[51] Millot P, San C, Bennana E, Porte B, Vignal N, Hugon J, Paquet C, Hosten B, Mouton-Liger F (2020) STAT3 inhibition protects against neuroinflammation and BACE1 upregulation induced by systemic inflammation. Immunology Letters 228, 129–134.

[52] Grandinetti KB, Jelinic P, DiMauro T, Pellegrino J, Fernández Rodríguez Rn, Finnerty PM, Ruoff R, Bardeesy N, Logan SK, David G (2009) Sin3B expression is required for cellular senescence and is up-regulated upon oncogenic stress. Cancer research 69, 6430–6437.

[53] Di Benedetto G, Iannucci LF, Surdo NC, Zanin S, Conca F, Grisan F, Gerbino A, Lefkimmiatis K (2021) Compartmentalized signaling in aging and neurodegeneration. Cells 10, 464.

[54] Lonze BE, Ginty DD (2002) Function and regulation of CREB family transcription factors in the nervous system. Neuron 35, 605–623.

[55] Myeku N, Clelland CL, Emrani S, Kukushkin NV, Yu WH, Goldberg AL, Duff KE (2016) Tau-driven 26S proteasome impairment and cognitive dysfunction can be prevented early in disease by activating cAMP-PKA signaling. Nature medicine 22, 46–53.

[56] Deyts C, Clutter M, Pierce N, Chakrabarty P, Ladd TB, Goddi A, Rosario AM, Cruz P, Vetrivel K, Wagner SL (2019) APP-mediated signaling prevents memory decline in Alzheimer’s disease mouse model. Cell reports 27, 1345–1355. e1346.

[57] Zlamal J, Althaus K, Jaffal H, Häberle H, Pelzl L, Singh A, Witzemann A, Weich K, Bitzer M, Malek N (2022) Upregulation of cAMP prevents antibody-mediated thrombus formation in COVID-19. Blood Advances 6, 248–258.

[58] Singh P, Gollapalli K, Mangiola S, Schranner D, Yusuf MA, Chamoli M, Shi SL, Lopes Bastos B, Nair T, Riermeier A (2023) Taurine deficiency as a driver of aging. Science 380, eabn9257.

[59] Rafiee Z, García-Serrano AM, Duarte JM (2022) Taurine supplementation as a neuroprotective strategy upon brain dysfunction in metabolic syndrome and diabetes. Nutrients 14, 1292.

[60] Thomas T, Stefanoni D, Reisz JA, Nemkov T, Bertolone L, Francis RO, Hudson KE, Zimring JC, Hansen KC, Hod EA (2020) COVID-19 infection alters kynurenine and fatty acid metabolism, correlating with IL-6 levels and renal status. JCI insight 5.

[61] Kim HY, Kim HV, Yoon JH, Kang BR, Cho SM, Lee S, Kim JY, Kim JW, Cho Y, Woo J (2014) Taurine in drinking water recovers learning and memory in the adult APP/PS1 mouse model of Alzheimer’s disease. Scientific reports 4, 7467.

[62] Baloni P, Funk CC, Yan J, Yurkovich JT, Kueider-Paisley A, Nho K, Heinken A, Jia W, Mahmoudiandehkordi S, Louie G (2020) Metabolic network analysis reveals altered bile acid synthesis and metabolism in Alzheimer’s disease. Cell Reports Medicine 1, 100138.

[63] Liu J, Li Z-B, Lu Q-Q, Yu Y, Zhang S-Q, Ke P-F, Zhang F, Li J-C (2022) Metabolite profile of COVID-19 revealed by UPLC-MS/MS-based widely targeted metabolomics. Frontiers in immunology, 3919.

[64] Stock AJ, Kasus-Jacobi A, Pereira HA (2018) The role of neutrophil granule proteins in neuroinflammation and Alzheimer’s disease. Journal of neuroinflammation 15, 1–15.

[65] Zenaro E, Pietronigro E, Bianca VD, Piacentino G, Marongiu L, Budui S, Turano E, Rossi B, Angiari S, Dusi S (2015) Neutrophils promote Alzheimer’s disease–like pathology and cognitive decline via LFA-1 integrin. Nature medicine 21, 880–886.

[66] Bawa KK, Krance SH, Herrmann N, Cogo-Moreira H, Ouk M, Yu D, Wu C-Y, Black SE, Lanctôt KL, Swardfager W (2020) A peripheral neutrophil-related inflammatory factor predicts a decline in executive function in mild Alzheimer’s disease. Journal of Neuroinflammation 17, 1–11.

[67] Cruz Hernández JC, Bracko O, Kersbergen CJ, Muse V, Haft-Javaherian M, Berg M, Park L, Vinarcsik LK, Ivasyk I, Rivera DA (2019) Neutrophil adhesion in brain capillaries reduces cortical blood flow and impairs memory function in Alzheimer’s disease mouse models. Nature neuroscience 22, 413–420.

[68] Vitte J, Michel BF, Bongrand P, Gastaut J-L (2004) Oxidative stress level in circulating neutrophils is linked to neurodegenerative diseases. Journal of clinical immunology 24, 683–692.

[69] Rosa BA, Ahmed M, Singh DK, Choreño-Parra JA, Cole J, Jiménez-Álvarez LA, Rodríguez-Reyna TS, Singh B, Gonzalez O, Carrion Jr R (2021) IFN signaling and neutrophil degranulation transcriptional signatures are induced during SARS-CoV-2 infection. Communications biology 4, 290.

[70] Akgun E, Tuzuner MB, Sahin B, Kilercik M, Kulah C, Cakiroglu HN, Serteser M, Unsal I, Baykal AT (2020) Proteins associated with neutrophil degranulation are upregulated in nasopharyngeal swabs from SARS-CoV-2 patients. PLoS One 15, e0240012.

[71] Bankar R, Suvarna K, Ghantasala S, Banerjee A, Biswas D, Choudhury M, Palanivel V, Salkar A, Verma A, Singh A (2021) Proteomic investigation reveals dominant alterations of neutrophil degranulation and mRNA translation pathways in patients with COVID-19. Iscience 24, 102135.

[72] Bathina S, Das UN (2015) Brain-derived neurotrophic factor and its clinical implications. Archives of Medical Science 11, 1164–1178.

[73] Lima Giacobbo B, Doorduin J, Klein HC, Dierckx RA, Bromberg E, de Vries EF (2019) Brain-derived neurotrophic factor in brain disorders: focus on neuroinflammation. Molecular neurobiology 56, 3295–3312.

[74] Maira D, Duca L, Busti F, Consonni D, Salvatici M, Vianello A, Milani A, Guzzardella A, Di Pierro E, Aliberti S (2022) The role of hypoxia and inflammation in the regulation of iron metabolism and erythropoiesis in COVID-19: The IRONCOVID study. American Journal of Hematology 97, 1404–1412.

[75] Demir B, Beyazyüz E, Beyazyüz M, Çelikkol A, Albayrak Y (2022) Long-lasting cognitive effects of COVID-19: is there a role of BDNF? European Archives of Psychiatry and Clinical Neuroscience, 1-9.

[76] Saiz-Sanchez D, Ubeda-Bañon I, Flores-Cuadrado A, Gonzalez-Rodriguez M, Villar-Conde S, Astillero-Lopez V, Martinez-Marcos A (2020) Somatostatin, olfaction, and neurodegeneration. Frontiers in Neuroscience, 96.

[77] Williams D, Yan BQ, Wang H, Negm L, Sackmann C, Verkuyl C, Rezai-Stevens V, Eid S, Vediya N, Sato C (2023) Somatostatin slows Aβ plaque deposition in aged APP NL-F/NL-F mice by blocking Aβ aggregation. Scientific Reports 13, 2337.

[78] Luty J, Hayward L, Jackson M, Duell PB (2021) Severe respiratory failure in a patient with COVID-19 and acromegaly: rapid improvement after adding octreotide. BMJ Case Reports CP 14, e243900.

[79] Chatterjee P, Pedrini S, Stoops E, Goozee K, Villemagne VL, Asih PR, Verberk IM, Dave P, Taddei K, Sohrabi HR (2021) Plasma glial fibrillary acidic protein is elevated in cognitively normal older adults at risk of Alzheimer’s disease. Translational psychiatry 11, 27.

[80] Cicognola C, Janelidze S, Hertze J, Zetterberg H, Blennow K, Mattsson-Carlgren N, Hansson O (2021) Plasma glial fibrillary acidic protein detects Alzheimer pathology and predicts future conversion to Alzheimer dementia in patients with mild cognitive impairment. Alzheimer’s Research & Therapy 13, 1–9.

[81] Aamodt AH, Høgestøl EA, Popperud TH, Holter JC, Dyrhol-Riise AM, Tonby K, Stiksrud B, Quist-Paulsen E, Berge T, Barratt-Due A (2021) Blood neurofilament light concentration at admittance: a potential prognostic marker in COVID-19. Journal of neurology, 1-10.

[82] Wang QL, Fang CL, Huang XY, Xue LL (2022) Research progress of the CXCR4 mechanism in Alzheimer’s disease. Ibrain 8, 3–14.

[83] Daoud S, Taha M (2022) Ligand-based Modeling of CXC Chemokine Receptor 4 and Identification of Inhibitors of Novel Chemotypes as Potential Leads towards New Anti-COVID-19 Treatments. Medicinal Chemistry 18, 871–883.

[84] Bezzi P, Domercq M, Brambilla L, Galli R, Schols D, De Clercq E, Vescovi A, Bagetta G, Kollias G, Meldolesi J (2001) CXCR4-activated astrocyte glutamate release via TNFα: amplification by microglia triggers neurotoxicity. Nature neuroscience 4, 702–710.

[85] Gavriel Y, Rabinovich-Nikitin I, Solomon B (2022) Inhibition of CXCR4/CXCL12 signaling: a translational perspective for Alzheimer’s disease treatment. Neural Regeneration Research 17, 108.

[86] Condrat CE, Thompson DC, Barbu MG, Bugnar OL, Boboc A, Cretoiu D, Suciu N, Cretoiu SM, Voinea SC (2020) miRNAs as biomarkers in disease: latest findings regarding their role in diagnosis and prognosis. Cells 9, 276.

[87] Kim Y-J, Kim SH, Park Y, Park J, Lee JH, Kim BC, Song WK (2020) miR-16-5p is upregulated by amyloid β deposition in Alzheimer’s disease models and induces neuronal cell apoptosis through direct targeting and suppression of BCL-2. Experimental Gerontology 136, 110954.

[88] Zhang N, Li W-W, Lv C-M, Gao Y-W, Liu X-L, Zhao L (2020) miR-16-5p and miR-19b-3p prevent amyloid β-induced injury by targeting BACE1 in SH-SY5Y cells. Neuroreport 31, 205–212.

[89] Li C, Wang R, Wu A, Yuan T, Song K, Bai Y, Liu X (2022) SARS-COV-2 as potential microRNA sponge in COVID-19 patients. BMC Medical Genomics 15, 1–10.

[90] Wicik Z, Eyileten C, Jakubik D, Simões SN, Martins Jr DC, Pavão R, Siller-Matula JM, Postula M (2020) ACE2 interaction networks in COVID-19: a physiological framework for prediction of outcome in patients with cardiovascular risk factors. Journal of clinical medicine 9, 3743.

[91] Ni W, Yang X, Yang D, Bao J, Li R, Xiao Y, Hou C, Wang H, Liu J, Yang D (2020) Role of angiotensin-converting enzyme 2 (ACE2) in COVID-19. Critical Care 24, 1–10.

[92] Frigerio CS, Lau P, Salta E, Tournoy J, Bossers K, Vandenberghe R, Wallin A, Bjerke M, Zetterberg H, Blennow K (2013) Reduced expression of hsa-miR-27a-3p in CSF of patients with Alzheimer disease. Neurology 81, 2103–2106.

[93] Harati R, Hammad S, Tlili A, Mahfood M, Mabondzo A, Hamoudi R (2022) miR-27a-3p regulates expression of intercellular junctions at the brain endothelium and controls the endothelial barrier permeability. PLoS One 17, e0262152.

[94] Zhou Y, Deng J, Chu X, Zhao Y, Guo Y (2019) Role of post-transcriptional control of calpain by miR-124-3p in the development of Alzheimer’s disease. Journal of Alzheimer’s Disease 67, 571–581.

[95] Kang Q, Xiang Y, Li D, Liang J, Zhang X, Zhou F, Qiao M, Nie Y, He Y, Cheng J (2017) MiR- 124-3p attenuates hyperphosphorylation of Tau protein-induced apoptosis via caveolin- 1-PI3K/Akt/GSK3β pathway in N2a/APP695swe cells. Oncotarget 8, 24314.

[96] Fernández-de Frutos M, Galán-Chilet I, Goedeke L, Kim B, Pardo-Marqués V, Pérez-García A, Herrero JI, Fernández-Hernando C, Kim J, Ramírez CM (2019) MicroRNA 7 impairs insulin signaling and regulates Aβ levels through posttranscriptional regulation of the insulin receptor substrate 2, insulin receptor, insulin-degrading enzyme, and liver X receptor pathway. Molecular and Cellular Biology 39, e00170–00119.

[97] Wei Z, Koya J, Reznik SE (2021) Insulin resistance exacerbates Alzheimer disease via multiple mechanisms. Frontiers in neuroscience 15, 687157.

[98] McDonald JT, Enguita FJ, Taylor D, Griffin RJ, Priebe W, Emmett MR, Sajadi MM, Harris AD, Clement J, Dybas JM (2021) Role of miR-2392 in driving SARS-CoV-2 infection. Cell reports 37, 109839.

[99] Wang Y, Zhu X, Jiang X-M, Guo J, Fu Z, Zhou Z, Yang P, Guo H, Guo X, Liang G (2021) Decreased inhibition of exosomal miRNAs on SARS-CoV-2 replication underlies poor outcomes in elderly people and diabetic patients. Signal Transduction and Targeted Therapy 6, 300.

[100] Lu T, Aron L, Zullo J, Pan Y, Kim H, Chen Y, Yang T-H, Kim H-M, Drake D, Liu XS (2014) REST and stress resistance in ageing and Alzheimer’s disease. Nature 507, 448–454.

[101] Hu G, Shi Z, Shao W, Xu B (2022) MicroRNA-214–5p involves in the protection effect of dexmedetomidine against neurological injury in Alzheimer’s disease via targeting the suppressor of zest 12. Brain research bulletin 178, 164–172.

[102] Fu XY (2006) STAT3 in immune responses and inflammatory bowel diseases. Cell research 16, 214–219.

[103] Mehla J, Singh I, Diwan D, Nelson JW, Lawrence M, Lee E, Bauer AQ, Holtzman DM, Zipfel GJ (2021) STAT3 inhibitor mitigates cerebral amyloid angiopathy and parenchymal amyloid plaques while improving cognitive functions and brain networks. Acta Neuropathologica Communications 9, 1–19.

[104] Matsuyama T, Kubli SP, Yoshinaga SK, Pfeffer K, Mak TW (2020) An aberrant STAT pathway is central to COVID-19. Cell Death & Differentiation 27, 3209–3225.

[105] Jafarzadeh A, Nemati M, Jafarzadeh S (2021) Contribution of STAT3 to the pathogenesis of COVID-19. Microbial pathogenesis 154, 104836.

